# The role of extra-striate areas in conscious motor behavior: a registered report with Fast-Optical Imaging

**DOI:** 10.1101/2024.04.23.590726

**Authors:** Elisabetta Colombari, Giorgia Parisi, Sonia Mele, Chiara Mazzi, Silvia Savazzi

## Abstract

Disclosing the brain areas responsible for the emergence of visual awareness and their timing of activation represents one of the major challenges in consciousness research. In particular, isolating the neural processes strictly related to consciousness from concurrent neural dynamics either related to prerequisites or post-perceptual processing has long engaged consciousness research. In this framework, the present study aims at unravelling the spatio-temporal dynamics underlying conscious vision by adopting a distinctive experimental design in which both awareness and motor response are manipulated, allowing the segregation of neural activity strictly related to awareness from response-related mechanisms. To this aim, we will employ a GO/NOGO detection task, in which participants will respond or withhold responding according to the experimental condition. Critically, during the performance of the task, participants’ brain activity will be recorded by means of Event-Related Optical Signal (EROS) technique, which provides accurate information about brain functions both from the temporal and spatial point of view, simultaneously. The combination of this experimental design with EROS recording will enable us to pinpoint the neural correlates underlying conscious vision and to disentangle them from processes related to the response. In addition, by coupling conventional EROS analysis with Granger Causality analysis, we will be able to clarify the potential interplay between consciousness-related extra-striate areas and response-related motor areas.

## 1. Introduction

Consciousness, namely the set of subjective experiences we have when we are awake, is one of the most intriguing topics debated in neuroscience research. In particular, the search for its neural correlates (NCC) has permeated the literature in recent decades. In broad strokes, one of the most widely used approaches to assess such NCCs involves contrasting brain activity occurring when a visual stimulus enters consciousness with brain activity occurring when the same stimulus does not reach awareness. This renowned paradigm is known as *contrastive analysis* (Aru *et al*., 2012) and has been frequently combined with electrophysiological recording or functional neuroimaging, leading to numerous and dissimilar results (Förster *et al*., 2020). Indeed, the interpretations of spatio-temporal dynamics underlying conscious vision are among the most disparate. ERP studies propose two possible electrophysiological markers as correlates of visual awareness: an earlier occipito-temporal negative deflection (i.e., Visual Awareness Negativity – VAN) detectable 200 ms after the presentation of the stimulus, and a later positivity (i.e., Late Positivity – LP) widespread over centro-parietal regions, peaking 300-500 ms after the stimulus onset (Koivisto & Revonsuo, 2010). However, the electrophysiological signature/s characterizing conscious vision has still to be elucidated. This may be attributed to one of the main limitations of the contrastive analysis, which is represented by its ineffectiveness in dissociating the true NCC (i.e., the set of neural correlates necessary and sufficient to enable consciousness) from concurrent neural dynamics either related to prerequisites or post-perceptual processing (Aru *et al*., 2012). In most prior studies aiming at identifying such NCCs, participants were asked to make judgments about their experience. However, such an operation could lead to confounding neural processes related to the task, not strictly to awareness per se.

For this reason, in an effort to disentangle the proper correlates of consciousness from neural activity related to the response, no-report paradigms have been employed. In this framework, no-report paradigms, where participants are not requested to perform any tasks or to provide any judgments about their perceptual experience, represent an advantageous tool to dissociate the neural processes strictly related to consciousness from subsequent processes related to the required response (Tsuchiya *et al*., 2015; Hatamimajoumerd *et al*., 2022). Studies employing this kind of paradigm with different techniques such as EEG and fMRI concluded that LP is highly modulated by several different cognitive processes occurring at later stages of processing (Mazzi *et al*., 2020; Schlossmacher *et al*., 2020; Dembski *et al*., 2021; Kronemer *et al*., 2022), as well as by the task relevance of the stimulus (Makeig & Jung, 2000; Pitts *et al*., 2014; Shafto & Pitts, 2015; Schelonka *et al*., 2017; Dellert *et al*., 2021; Hense *et al*., 2024). By contrast, the role of response requirements, as well as that of attention, on the VAN are still debated as different studies have reported both positive (e.g., Bola & Doradzińska, 2021; Dellert *et al*., 2021; Doradzińska & Bola, 2024) and negative (e.g., Koivisto *et al*., 2006; Cohen *et al*., 2020; Dellert *et al*., 2022; Ciupińska *et al*., 2024) results. Interestingly, in a study published in 2016 by Koivisto and colleagues (Koivisto *et al*., 2016), authors successfully dissociated ERP correlates of visual awareness from those related to post-perceptual mechanisms, disclosing that VAN was not modulated by response requirements. The authors adopted a particular partial-report paradigm in which participants were sometimes asked to provide a report by pressing a response button when they were aware of the stimulus and sometimes to withhold responding in case of awareness. They found that, while the amplitude of LP was modulated by the response (i.e., it was greater in trials where participants were asked to respond in case of awareness, compared to the Aware condition where they were asked to withhold responding), VAN did not change depending on task requirements. This allowed Koivisto and colleagues to advocate for an early onset of visual awareness: the phenomenal content of a visual experience, indeed, takes place before LP, more specifically in the temporal window of VAN.

Several pieces of evidence are consistent in considering VAN as the electrophysiological signature of phenomenal consciousness (Koivisto *et al*., 2008; Railo *et al*., 2015), while the localization of its neural generator still remains open. In this regard, previous MEG source localization studies (Vanni *et al*., 1996; Liu *et al*., 2012) identified the Lateral Occipital Complex (LOC), an extra-striate visual areas traditionally associated with objects recognition, as the generator of VAN. Moreover, previous no-report studies using both EEG and fMRI (Dellert et al., 2021; Kronemer et al., 2022) have also found awareness effects in LOC and linked it to VAN. The same result was achieved in a recent work aimed at unravelling the spatio-temporal dynamics underlying conscious vision (Colombari *et al*., 2024). In such study, participants were asked to perform a discrimination task on the orientation of a tilted Gabor patch while their brain activity was recorded first with EEG and then with Fast Optical Imaging. This allowed authors to identify the exact temporal window of VAN and LP and then, by taking advantage of the peculiarity of Fast Optical Imaging of achieving both temporal and spatial accurate information (Gratton & Corballis, 1995; Gratton & Fabiani, 2010; Baniqued et al., 2013), to investigate the spatio-temporal unfolding of brain activity occurring in these predetermined time windows. Authors contrasted activity of Aware trials (i.e., trials in which participants reported to perceive the orientation of the stimulus) with activity of Unaware ones and observed a sustained activation of LOC in the VAN temporal window, consistently with the above-mentioned MEG studies. More interestingly, they observed that, only when the stimulus crossed the threshold of consciousness, activity in extra-striate visual areas triggered subsequent activation of motor areas, although motor response was required in both Aware and Unaware conditions. Authors tried to interpret this unexpected finding by ascribing it to the selection of the correct response, that could be provided in the Aware trials only where participants consciously perceived the stimulus. Indeed, in Aware trials participants had to press a specific button on the response box (to provide the correct answer about the orientation of the Gabor patch), while when the stimulus was unseen (i.e., Unaware trials) they had to respond randomly, by pressing indifferently one of the two response buttons. However, the employed experimental paradigm did not allow the authors to thoroughly investigate this issue. Thus, in order to clarify the interplay between extra-striate areas and motor regions in awareness, in the present study we will adopt a go/no-go detection task (similar to that adopted by Koivisto *et al*., 2016), while recording participants’ brain activity by means of Fast Optical Imaging. Specifically, Event-Related Optical Signal (EROS) technique will be employed. This technique, by shedding near-infrared light through the brain tissues, is able to detect changes in the light scattering properties that are known to be directly related to neural activity, thus providing accurate information about brain functions both from the temporal and spatial point of view, simultaneously (Gratton *et al*., 1997; Gratton & Fabiani, 1998, 2001). Critically, the study will adopt a distinctive paradigm manipulating both awareness and response. The latter, indeed, will be provided sometimes in the Aware condition (condition Aware-GO/Unaware-NOGO) and sometimes in the Unaware one (condition Aware-NOGO/Unaware-GO). This double manipulation will enable us to unravel the spatio-temporal unfolding of awareness-related activity, by disentangling neural activity related to awareness from response-related mechanisms. Indeed, in the present study, we can investigate the NCCs both when the motor response is required and when no task is performed, thus allowing to isolate consciousness effects from the effects related to the task. Importantly, the experimental paradigm adopted will enable us to elucidate the interplay between extra-striate visual areas and motor areas. Indeed, in addition to conventional EROS analyses, we will perform Granger Causality analysis, in order to disclose the relationship existing among the investigated areas. In broad strokes, Granger analysis allows to move beyond the classical identification of cortical activation provided by EROS analysis by disclosing functional circuits underpinning the investigated brain function (Seth *et al*., 2015). When coupled with EROS, Granger Causality analysis represents a powerful tool to highlight predictive relationship between activations in the investigated regions of interest (ROI) at different time-points (Parisi *et al*., 2020).

Based on previous literature suggesting that VAN is independent from subjective report (Koivisto *et al*., 2016; Ye *et al*., 2024) and LOC represents the cortical generator or VAN (Liu *et al*., 2012; Colombari *et al*., 2024), we expect Aware trials to elicit early greater activation of LOC, independently of the response requirement. Moreover, by combining EROS conventional analysis with Granger Causality analysis, and manipulating both awareness and motor response, we aim to highlight potential interplay between consciousness-related extra-striate areas and response-related motor areas both when the motor response is required and when it has to be inhibited.

## 2. Methods

### 2.1 Ethics Information

The study is approved by the local Ethics Committee (Prog.171CESC) and it will be conducted in accordance with the principles laid down in the 2013 Declaration of Helsinki. Participants will be recruited from the University of Verona community, by means of printed flyers displayed on notice boards at different University of Verona sites and through advertisements on social media. Each participant will be fully informed about the modalities of the study before taking part in the experiment and written informed consent will be signed. In addition, participants will receive compensation for their participation and will be debriefed after the conclusion of the experiment.

### 2.2 Participants

We will recruit healthy adults, right-handed (as assessed by means of the standard handedness inventory *Edinburgh Handedness Questionnaire*; Oldfield, 1971) and aged between 18 and 50 years old. All of them will have to report normal or corrected-to-normal vision, no history of neurological or psychiatric disorders and no contraindications to MRI. The study will be conducted at the PandA lab of the University of Verona (Italy).

#### 2.2.1 Sample size estimation

The estimate of the sample size for the current study is based on our previous EROS study (Colombari et al., 2024), in which a similar paradigm was adopted, and similar analyses on similar ROIs were performed. Specifically, EROS data from the ROI of LOC were extracted, and significant time-points were averaged within participants so to have one value for each of them. Then, a one-sample t-test was performed (t(23) = 2.99, p = .006, Cohen’s d = .611), and the resulting Cohen’s d was employed to compute the sample size estimation for the current study. Specifically, the estimated sample size for research questions Q1 (i.e., “Can we replicate Colombari et al., 2024 findings showing that LOC is an NCC?”) and Q2 (i.e., “Is the activity in LOC independent from the response?”) was calculated with G-Power software (v. 3.1.9.7), with a power of 90% and a level of significance of 2%. The estimated sample size resulted in 32 participants (critical t= 2.143; actual power= 0.900). Considering that the estimated sample size for this study (n=32) is more than double the typical sample size of EROS studies present in literature, the same estimated sample size seems to be also adequate to answer research questions Q3 (i.e., “Does consciousness modulate the activation of motor areas in a detection task?”) and Q4 (i.e., “Does consciousness modulate the activation of motor areas in ABSENCE of motor response?”). For a review of the existing EROS literature, see Supplementary Table 1 at https://osf.io/ebfu3/?view_only=9ec2e6bf32ba4a8bb8b858639ec40a59) from which emerges that, on average, EROS studies employ experimental samples composed of about 13 participants (mean 12.944; SD 7.008).

### 2.2.2 Exclusion Criteria

As better specified in section 2.3, before getting involved in the study, participants will undergo a perceptual threshold assessment, in order to identify the proper stimulus to be employed in the main experiment. To be enrolled in the study, participants will have to successfully complete this session. The criterion used is that one of the stimuli presented during the threshold assessment will have to be acknowledged as perceived a minimum of 25%, a maximum of 75%, or closest to the 50% of the times (i.e., at perceptual threshold level). If no stimulus results at the threshold level, the participant should not be enrolled in the study.

In addition, participants who will not complete all the experimental sessions, as well as participants reporting a level of Awareness superior to 75% or inferior to 25% at the end of the experiment will be excluded from analyses. This is to maintain comparable the number of trials in the two experimental conditions (i.e., Aware and Unaware) and to ensure a reliable EROS activity (because of its relatively low signal-to-noise ratio, EROS needs a high number of trials per condition, in order to compute statistics). Moreover, participants whose behavioral performance will be affected by biases related to the behavioral response (as assessed by catch trial analysis, explained more in detail below) will be excluded from the analyses (see below –*Section 2.8.1 Behavioral data* for more detailed information on the analysis of catch trials). Finally, participants whose EROS signal could not be detected properly during the experiment (for example because of too dark hair or technical issues) will not be included in the analyses as well. In particular, the opacity value (i.e., the product of the scattering and absorption coefficients) will be estimated for each participant. Based on this value, it is possible to judge the quality of the signal for each participant, independently from the experimental condition. Opacity values of all participants will be averaged together providing the absorption coefficient to be used when running statistical analysis.

Participants whose opacity value is equal to 0 or exceeds three standard deviations of the mean will be excluded from statistical analyses.

Importantly, each participant who will be excluded due to the previously mentioned exclusion criteria, will be replaced with the recruitment of another participant. Thus, the number of participants to be recruited will be increased to reach a total of 32 analyzed subjects, as specified in section 2.2.1.

### 2.3 Stimuli

Stimuli will be created by means of a custom-made Matlab script (version R2022b; the MathWorks, Inc., Natick, MA) and resized by means of Photoshop (Adobe Photoshop CC, v2014.0.0). As shown in Figure 1, they will be gray circles (.85 .85 .85 RGB), presented on a black background, with 8 radii equally distanced one from another. One radius (the first one, clockwise) can be slightly thicker than the others (critical trials) or not (catch trials). The thickness of the radius for critical stimuli will be individually assessed for each participant on the basis of a subjective perceptual threshold assessment that will be held before the main experiment.

**Figure 1.**
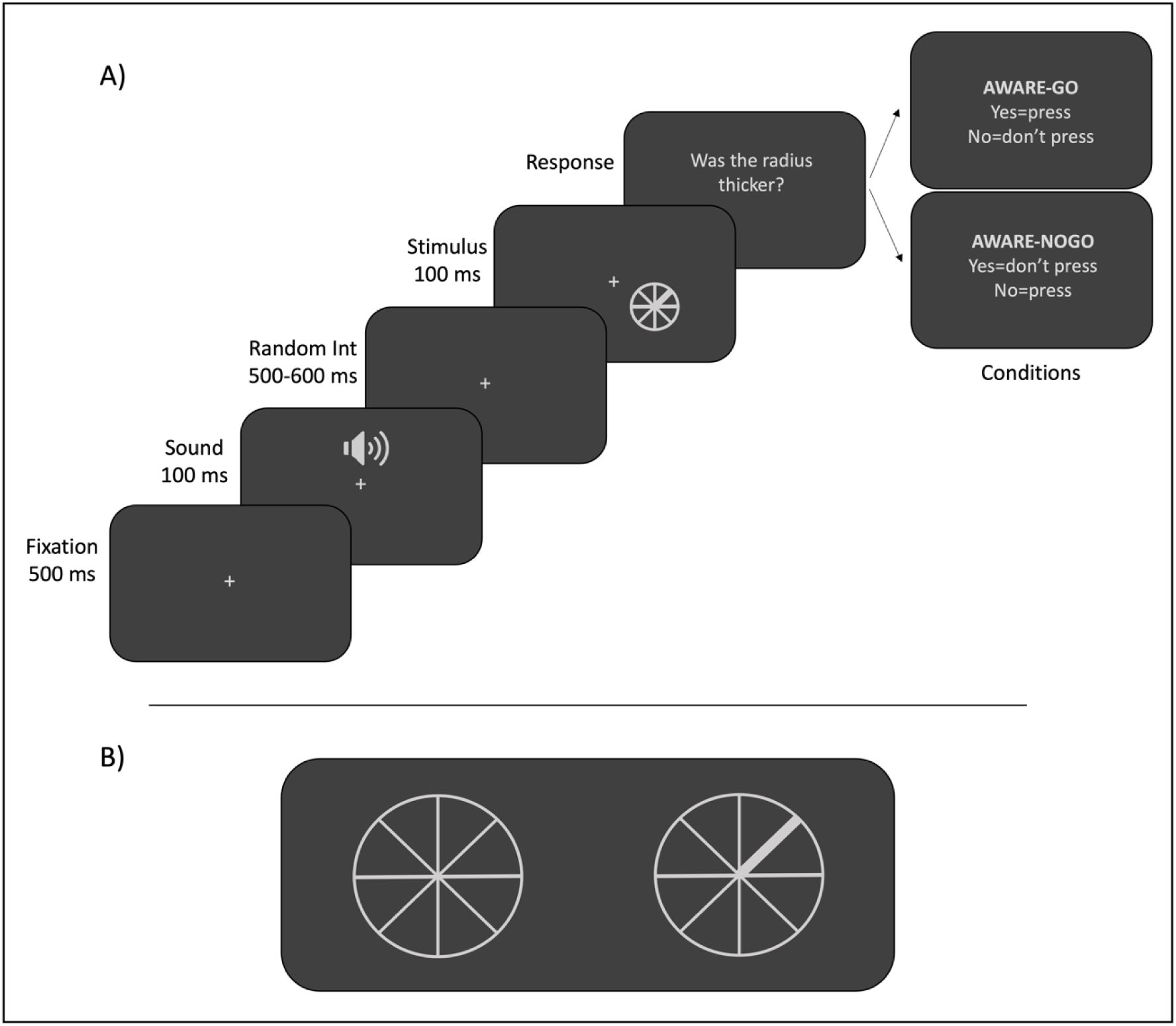
Trial procedure and stimuli: **A)** Experimental procedure: the trial begins with a fixation cross persisting at the center of the screen for 500 ms. After that, an acoustic tone lasting 100 ms will be presented, followed by a random interval ranging from 500 to 600 ms. Then, the stimulus will be presented for 100 ms and participants will be asked to respond according to the experimental condition (i.e., Aware-GO or Aware-NOGO). **B)** Example of stimuli: on the left is shown the catch stimulus, with all the radii equally thick; on the right is depicted the critical stimulus, with the first radius, clockwise, thicker than the others.

Both in the perceptual threshold assessment and in the main experiment, the stimulus will be presented in the lower right quadrant of the screen, specifically at an eccentricity of 3.5° from the fixation cross along the vertical meridian and of 2° along the horizontal one. This is to allow a left-lateralized EROS montage, as a full-head montage is not achievable in our lab due to technical constraints (i.e., insufficient probes). Moreover, since EROS technique is sensitive to depth, a right-lateralized stimulus ensures that it elicits activity in the left portion of the primary visual cortex, which is known to be anatomically closer to the skull compared to the right one (Zhao *et al*., 2022), thus ensuring a better penetration of near-infrared light through brain tissues.

### 2.4 Perceptual Threshold Assessment

Before starting the experiment, participants will undergo a perceptual threshold assessment, with the aim of identifying, for each participant, the level of thickness of the critical radius so that it results to be perceived as thicker 50% of the times. To this aim, stimuli with different levels of radius thickness will be randomly presented and the subjective perceptual threshold will be measured using the method of constant stimuli. Specifically, 9 levels of radius thickness will be presented. The range of stimuli to be used in the perceptual threshold assessment will be selected based on the results of a pilot experiment in which participants were asked to perform the same task employed in the perceptual threshold assessment while presented with a wider range of radius thickness. This will allow us to identify a smaller range of optimal stimuli to be presented thus excluding a range of stimuli whose thickness was almost never or always reported by participants. Each level of radius thickness will be presented 5 times per block, for a total of 8 blocks. Thus, all the stimuli, as well as the catch stimulus, will be presented 40 times each. Participants will be asked to press the spacebar as soon as they detect the stimulus with a thicker radius. The stimulus identified as perceived a minimum of 25%, a maximum of 75%, and closest to 50% of the times at the end of the subjective perceptual threshold assessment will be used in the experimental task, together with the catch. The perceptual threshold assessment, as well as the main experiment, will be conducted in a dimly illuminated room and participants will be sitting in front of a 17 in. LCD monitor (resolution 1920×1080, refresh rate of 144 Hz) placed at a viewing distance of 57 cm. Their head will be held in place by means of an adaptable chin rest so that eyes are aligned with the center of the screen. Both the perceptual threshold assessment and the main experiment will be programmed and administered using E-Prime 3.0 software (E-Prime Psychology Software Tools Inc., Pittsburgh, PA, USA). Before starting the perceptual threshold assessment, participants will undergo a fixation training (Leung *et al*., 2009), in order to ensure they will maintain their gaze on the central fixation cross correctly.

### 2.5 Experimental Procedure

The experiment will be composed of two identical sessions lasting approximately 3 hours each performed on different days. The first session will be preceded by the assessment of the subjective perceptual threshold, which, in turn, will last around 20 minutes. The two experimental sessions will be identical except for the EROS montages, specifically devised to obtain better coverage of the brain areas of interest. The order of the montages will be counterbalanced across participants, as well as the order of conditions (see below for more detailed information).

The task will be a two-conditions go/no-go detection task, similar to that adopted by Koivisto *et al*., 2016, in which participants have to respond in different ways according to the experimental condition (Table 1). In condition “Aware-GO”, they will be asked to press the spacebar on the keyboard as soon as they perceive the thicker radius, and withhold responding when they do not perceive any difference among radii. On the contrary, in condition “Aware-NOGO”, participants will be asked to withhold responding when they perceive a thicker radius, and press the response button when they do not perceive any difference. Each trial will begin with the presentation of a central fixation cross, followed 500 ms later by a sound (1000Hz) presented for 100 ms, notifying participants of the subsequent onset of the stimulus. After a random interval ranging from 500 to 600 ms, the stimulus will be presented for 100 ms in the lower right quadrant of the screen. After that, participants will be asked to respond according to the experimental condition. Each experimental session will be composed of 24 blocks: 12 blocks for condition Aware-GO/Unaware-NOGO and 12 blocks for condition Aware-NOGO/Unaware-GO, counterbalanced across participants according to the order depicted in Table 1. Each block will consist of 50 critical trials and 15 catch trials. The whole experiment will be composed of 48 blocks per participant, for a total of 2400 critical trials and 720 catch trials per participant.

**Table 1.** Experimental conditions. Both Awareness and Response are manipulated: Awareness is experimentally manipulated by employing a threshold stimulus, so that sometimes it is consciously perceived (Aware) and sometimes not (Unaware). Response is manipulated by the task: in condition GO participants are asked to respond by pressing a key, while in condition NOGO they are asked to withhold responding. The combination of these two manipulations gives rise to the 4 experimental conditions depicted in the table.

|  |  | Awareness |  |
| --- | --- | --- | --- |
|  |  | yes | no |
| Response | no | Aware-NOGO | Unaware-NOGO |
|  | yes | Aware-GO | Unaware-GO |

**Table 2.**
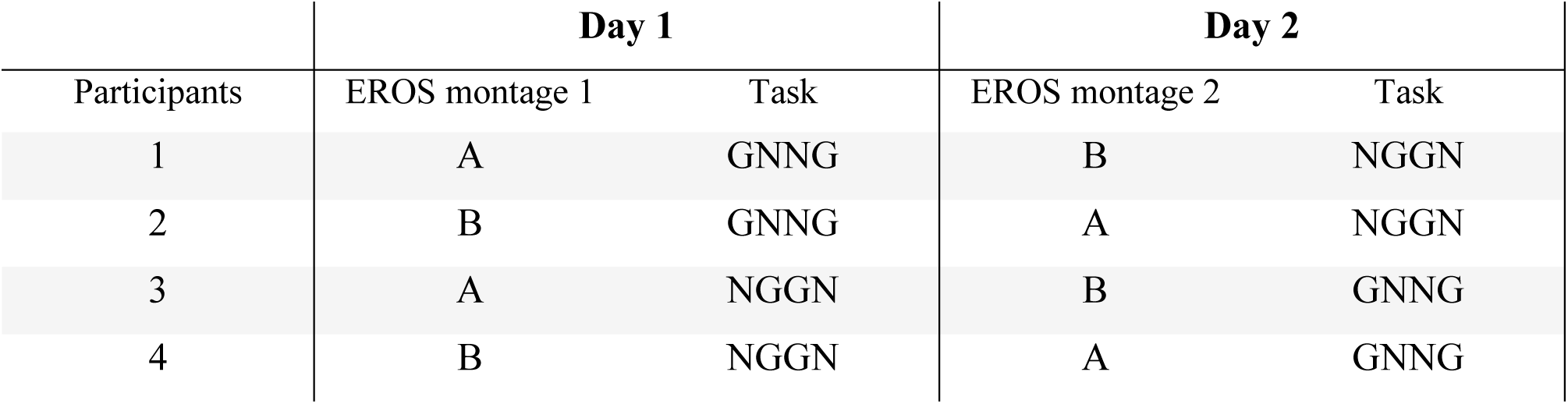
Counterbalancing of montages and task conditions across participants. Both EROS montages and task conditions (G = Aware-GO/Unaware-NOGO; N = Aware-NOGO/Unaware-GO) will be counterbalanced across participants. In the column “Task”, each letter represents 6 blocks of task. Thus, each day, participants will perform 12 blocks per condition, for a total of 24 blocks of task per day.

| Participants | Day 1 |  | Day 2 |  |
| --- | --- | --- | --- | --- |
|  | EROS montage 1 | Task | EROS montage 2 | Task |
| 1 | A | GNNG | B | NGGN |
| 2 | B | GNNG | A | NGGN |
| 3 | A | NGGN | B | GNNG |
| 4 | B | NGGN | A | GNNG |

### 2.6 Optical Recording

Three synchronized Imagent frequency domain systems (ISS, Inc., Champaign, IL) will be used to record continuous fast optical data throughout experimental sessions. Each system is equipped with 4 photo-multiplier tubes detectors, for a total of 12 detectors. Near-infrared light (830 nm) will be delivered from 48 laser diodes on participants’ scalp and it will be modulated at 110 MHz. Each of 12 detectors will receive light from sets of 16 light emitters, multiplexed every 25.6 ms, resulting in a sampling rate of 39.0625 Hz.

To avoid cross-talk between channels, the array of source-detector pairs (i.e., the montage) will be created by means of a specific program (NOMAD, Near-Infrared Optode Montage Automated Design) implemented in Matlab, useful to place sources and detectors at optimal distances. In this experiment, we will set the minimal distance to 17.5 mm and the maximum distance to 50 mm, in order to ensure an extensive coverage of the brain regions of interest both from the spatial and the depth point of view. The distance between the source and the detector of a channel, in fact, determines the depth of the light pathway (Gratton *et al*., 2000), thus corresponding to the depth of the investigation: namely, longer channels can investigate deeper layers and shorter channels can examine shallower regions.

Both light emitters and detectors will be placed on participants head using a custom-built helmet. To minimize interferences, before placing the optical fibers on the head, the hair will be carefully moved with cotton buds, so that the fibers can reach the scalp directly. In order to better adhere to the head of the participant, we will employ two helmets of different sizes: one 55-56 cm large, and one 57-58 cm large. For each helmet, we will develop two different montages, so that to provide a dense coverage of the regions of interest (i.e., the left occipital, temporal and parietal cortices, see Figure 2). Each montage will consist of the combination of 12 detectors and 48 light emitters, resulting in a total of 192 channels per montage. As mentioned before, each montage will be recorded in a separate session, and the order will be counterbalanced across participants.

**Figure 2.**
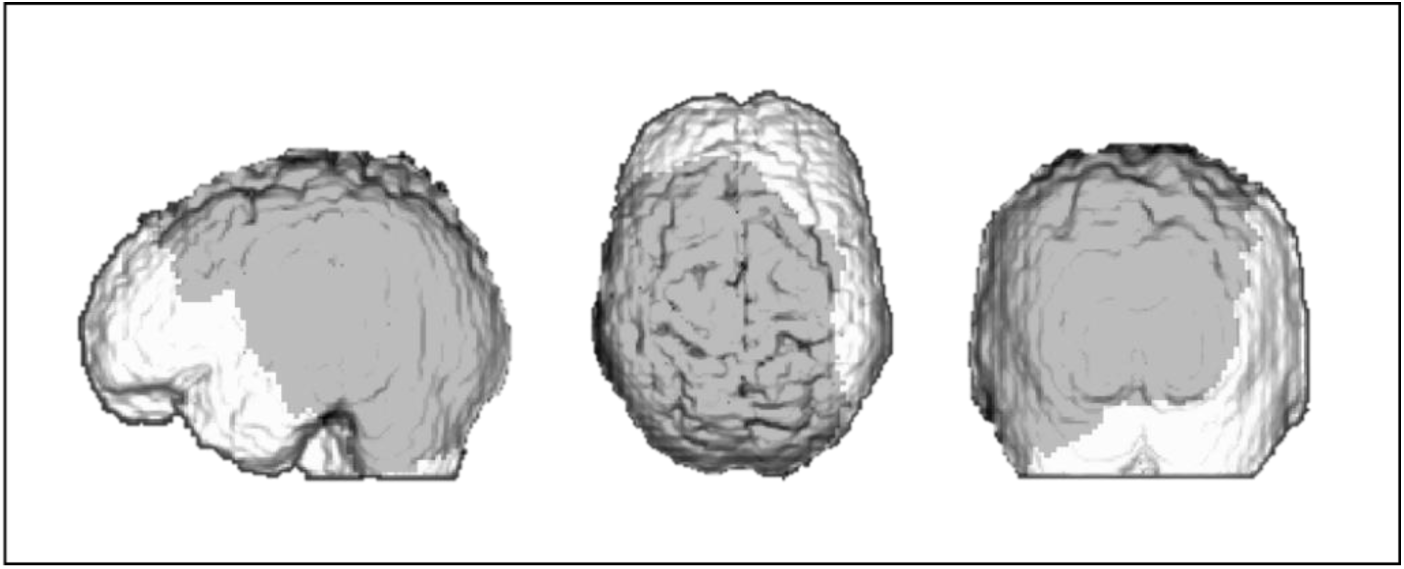
Covered area. The gray area represents the area covered by the EROS montages (combined together) from the sagittal, axial and coronal point of view.

At the end of each EROS session, the scalp location of each source and detector will be digitized in relation to four fiducial points (i.e., nasion, inion and pre-auricular points) with a neuro-navigation software (SofTaxic, E.M.S., Bologna, Italy) combined with a 3D optical digitizer (Polaris Vicra, NDI, Waterloo, Canada). Afterwards, the digitized scalp locations will be co-registered with each participant’s individual MRI, using a dedicated software package (OCP, Optimized Co-registration Package, Matlab code developed by Chiarelli and colleagues (Chiarelli *et al*., 2015).

For this reason, participants will undergo a structural MRI at the Azienda Ospedaliera Universitaria Integrata of Verona (AOUI).

### 2.7 MRI Acquisition

Participants’ individual structural MRI will be acquired by means of a 3 Tesla Philips Ingenia scanner with a 32-channel head RF receive coils. A whole brain high-resolution 3D T1-weighted image (T1w) Turbo-field echo image (1mm-isotropic TE/TR=3.8/8.4 ms, TI=1050 ms) will be acquired.

The T1w field of view (240 x 240 x 180 mm) will be large enough to allow for the ears and the entire scalp to be fully included in the image to facilitate later and accurate co-registration with functional data.

### 2.8 Data Analysis

#### 2.8.1 Behavioral data

Raw data will be processed by means of custom scripts created on Matlab (the MathWorks, Inc., Natick, MA). Data will be divided into the 4 experimental conditions (i.e., Aware-GO, Unaware-NOGO, Aware-NOGO, Unaware-GO). For each participant, trials with reaction times lower than 150 ms and higher than 3 standard deviations from the mean will be excluded from the analysis. Data will be successively analyzed using Jamovi (version 2.3.28): first, the percentage of Aware and Unaware trials will be calculated, in order to assess that a sufficient amount of trials is present for each condition. Participants presenting more than 75% or less than 25% of Awareness will be discarded from the sample. This is because, in that case, the number of Unaware (or Aware) trials would be insufficient for statistical EROS analysis. EROS technique, indeed, although having a high localization power from both the spatial and temporal point of view, has a relatively low signal-to-noise ratio. For this reason, a high number of trials is required for statistical analysis. Subsequently, reaction times (RTs) will be analyzed for the “GO” conditions, thus paired sample t-tests (two-tailed) will be applied to compare the mean RTs between Aware-GO and Unaware-GO conditions. Finally, to verify that participants are performing the task accurately and that there are no biases related to the response, catch trials will be analyzed. As mentioned above, catch trials are those trials in which all the radii of the stimulus are equally thick, thus no differences in the stimulus are present. In case of catch trials, the participants’ task will be different according to the condition: in the Aware-GO condition, they are expected to withhold responding, while in the Aware-NOGO condition, they are expected to respond. Thus, catch trials will be analyzed separately for the two conditions (GO and NOGO) by means of a paired sample t-test (two-tailed), in order to ensure that the behavioral performance follows the above-mentioned trend. Paired sample t-tests (two-tailed) will indeed be performed to test whether catch trials performance is significantly different from critical trials.

#### 2.8.2 EROS data

Pre-processing of continuous phase delay (i.e., time-of-flight) data will be computed by means of a dedicated in-house software, P-POD (Pre-Processing of Optical Data, run in Matlab, version R2013b). Thus, raw data will be normalized (i.e., corrected for phase wrapping and de-trended to remove low-frequency drifts), demeaned and filtered by means of a 6^th^ order Butterworth band-pass filter which allows frequencies between 0.5 Hz and 15 Hz. Pulse artifact will be removed by using a regression algorithm (GRATTON *et al*., 1995). After that, data will be averaged separately for each subject, condition, and channel and segmented into epochs time-locked to the onset of the stimulus. Each epoch will comprise a period from 486 ms before the stimulus onset to 998 ms following the stimulus onset, resulting in an epoch lasting 1484 ms. Subsequently, statistical analyses will be computed with an in-house software package (Opt-3d; (Gratton, 2000)), which provides statistical spatial maps of fast optical data.

To perform statistics, data from channels whose diffusion paths intersect a given voxel will be combined (Wolf *et al*., 2014). Phase delay data will be spatially filtered with an 8-mm Gaussian kernel and baseline corrected using a 204 ms time-window preceding the stimulus onset. Within each ROI, t-Statistics will be calculated at group level, converted into Z-scores and corrected for multiple comparisons using random field theory (Worsley *et al*., 1995; Kiebel *et al*., 1999). Then, Z-scores will be weighted and orthogonally projected onto the surface of an MNI template brain, according to the physical homogenous model (Arridge & Schweiger, 1995; Gratton, 2000).

In order to investigate the neural dynamics related to conscious vision and to disentangle the role of the motor areas, the following contrasts between conditions will be computed: 1) Aware-GO versus Unaware-GO and 2) Aware-NOGO versus Unaware-NOGO. These contrasts allow to investigate the research questions the proposed study aims at answering (see Section 3 for a detailed description of the planned analysis). Importantly, both frequentist and Bayesian statistics (with default priors) will be computed, to test both positive and negative effects.

Moreover, Granger Causality analysis will be computed. Granger Causality analysis allows to explore the predictive interactions between different brain areas at different time-points. Specifically, this approach requires a region of interest (ROI) to be used as a “seed” and investigating whether the activity of this seed predicts activity in the other ROIs at a later time-lag, by deriving statistical maps from t-statistics computation (then transformed into z scores) for each lag.

Statistical functional analysis will be computed within specific predetermined regions of interest (ROIs) and time intervals. ROIs will be defined by a 2-dimensional box-shaped structure, covering an area of 20×20 millimeters. Critical ROIs will be selected on the basis of the results obtained in the above-mentioned experiment (Colombari *et al*., 2024) and they will be located in the occipital and in the left parietal and temporal lobes, specifically over the primary visual cortex (V1, Brodmann Area 17), the left lateral occipital cortex (LOC, Brodmann Area 19), the left supplementary motor area (SMA, Brodmann Area 6), the left premotor area (PM, Brodmann Area 6) and the left primary motor cortex (M1, Brodmann Area 4). Statistical analysis will be computed within specific temporal windows of interest selected on the basis of the results obtained by Colombari et al., 2024. This is to reduce the risk of false positives, as Opt3d does not offer the possibility to correct data for multiple comparisons in the temporal domain. The specific time windows tested for each hypothesis are listed in Table 3.

## 3. Study design

| Question | Hypothesis | Sampling Plan | Analysis Plan | Rationale for deciding the sensitivity of the test for confirming or disconfirming the hp | Interpretation given different outcomes |
| --- | --- | --- | --- | --- | --- |
| **Q1:** Can we replicate Colombari et al., 2024 findings showing that LOC is an NCC? | **H1:** We hypothesize to replicate Colombari et al., 2024 results: greater activity in LOC in an early temporal window (i.e., 150-350 ms post stimulus onset) is observed when contrasting Aware and Unaware trials in the condition in which the response is required (i.e., GO condition)<br>**Expected outcome:**<br>LOC aware-GO>LOC unaware-GO, as measured by EROS activity | Since the present research question aims at replicating the results of Colombari et al., 2024, sample size estimation is based on those EROS data. Sample size calculation was ~~thus~~ performed with G-Power software (v. 3.1.9.7), with a power of 90% and a significance level of 2%, resulting in 32 participants. | **A1:** The goal is to replicate the results of Colombari et al., 2024, in which the manual response was required for both Aware and Unaware conditions. Here, in order to perform the same analysis, early LOC activity in Aware-GO and Unaware-GO trials will be compared by using a paired-sample one-tailed t-test, computed with the EROS dedicated analysis software “Opt3d”.<br>In addition to frequentist statistics, Bayesian statistics (with default prior) will be computed to test potential null effects.<br>**Contrast to be computed:**<br>AWARE GO VS<br>UNAWARE GO<br>**ROI to be tested:**<br>LOC | Sample size estimation is based on previous EROS data (Colombari et al., 2024), on which similar analyses were performed. | **O1.1:** A significant t-test within the interval of interest will be interpreted as a successful replication of previous findings, supporting the involvement of LOC in NCC.<br>**O1.2:** The absence of this effect will not confirm the hypothesis, suggesting that LOC is not involved in the conscious detection of a stimulus property. |
|  |  |  | **Time interval of interest:**<br>150-350ms after stimulus onset |  |  |
| **Q2:** Is the activity in LOC independent from the response? | **H2:** We hypothesize that LOC activity is independent from response requirement: when contrasting activity elicited by Aware-NOGO trials with activity elicited by Unaware-NOGO trials, we expect to find the same activation of LOC found in the Aware-GO vs Unaware-GO contrast.<br>**Expected outcome:**<br>LOC aware-NOGO>LOC unaware-NOGO, as measured by EROS activity<br>(LOC aware-GO>LOC unaware-GO)=(LOC aware-NOGO>LOC unaware-NOGO) | Since research question Q2 involves the same analyses of research question Q1 (but for the NOGO condition), the sampling plan for Q2 is the same as for Q1. | **A2.1:** A paired-sample one-tailed t-test will be computed in order to compare early activity in LOC in the NOGO condition. Thus, activity in Aware-NOGO and Unaware-NOGO trials will be contrasted.<br>Both frequentist and Bayesian statistics (with default prior) will be computed.<br>**Contrast to be computed:**<br>AWARE NOGO VS UNAWARE NOGO<br>**ROI to be tested:**<br>LOC<br>**Time interval of interest:**<br>150-350ms after stimulus onset | As above | **O2.1.1:** A significant t-test in the time window of interest will suggest that LOC activity is independent from response, since its activity is observed even when no response is required (NOGO conditions).<br>**O2.1.2:** If greater activity in LOC in the time window of interest is not observed, then it means that LOC activity is somehow related to the motor response. |
|  |  |  | **A2.2:** The interaction effect between awareness and motor response will be tested by means of a paired-sample one-tailed |  | **O2.2.1:** Significant interaction effect will suggest that activity in LOC depends from response requirement<br>**O2.2.2:** The absence of a difference between the two effects will suggest |
|  |  |  | t-test computed between contrast Aware-GO VS Unaware-GO and contrast Aware-NOGO VS Unaware-NOGO<br>In addition to frequentist statistics, Bayesian statistics (with default prior) will be computed to test potential null effects<br>**Contrast to be computed:**<br>(AWARE GO VS UNAWARE GO) - (AWARE NOGO VS UNAWARE NOGO)<br>**ROI to be tested:**<br>LOC<br>**Time interval of interest:**<br>150-350ms after stimulus onset |  | that motor response does not affect awareness-related activity in LOC |
| **Q3:** Does consciousness modulate activation of motor areas in a detection task? | **H3:** When a motor response is required, consciousness modulates activation of motor areas (MA), as activity in motor areas is triggered by LOC (Colombari et al., 2024)<br>**Expected outcome:** | Considering that the estimated sample size for this study (n= 32) is more than twice the typical sample size of EROS studies present in literature (see Supplementary Table 1, where a systematic review of existing EROS literature revealing that the typical sample size used is | **A3.1** A paired-sample one-tailed t-test will be computed in order to compare early activity in Motor Areas in the GO condition. Thus, activity in Aware-GO and Unaware-GO trials will be contrasted.<br>In addition to frequentist statistics, Bayesian | As above | **O3.1.1:** A statistically significant difference between the two conditions will suggest that, even in a detection task, response related motor activity is stronger in the Aware condition compared to the Unaware one.<br>In Colombari et al., 2024 this difference was observed. Importantly, in this previous study a *discrimination* task was employed and participants were asked to provide two different |
|  | MA aware-GO>MA unaware-GO, as measured by EROS activity<br>LOC activity predicts MA activity (investigated by means of Granger Causality Analysis) | 13 participants is depicted), the same estimated sample size of Q1 and Q2 seems to be also adequate to answer research questions Q3 and Q4. | statistics (with default prior) will be computed to test potential null effects<br>**Contrast to be computed:**<br>AWARE GO VS UNAWARE GO<br>**ROI to be tested:**<br>Motor areas<br>**Time interval of interest:**<br>Based on mean RTs, with a time window of $\pm 1.5$ sd around the mean |  | responses in case of Awareness (intentional) or Unawareness (random). Instead, in this study participants are asked to perform a *detection* task, in which the motor behavior made to provide the response, when required, is the same for both Aware and Unaware condition and thus no response selection is required.<br>**O3.1.2:** If no difference between the tested conditions is observed, it will suggest that in a detection task there is no difference in the motor activity related to the response. |
|  |  |  | **A3.2:** In order to further investigate the flow of activity occurring in the investigated brain areas, Granger Causality Analysis will be performed. In the present study, we will perform Granger analysis on the “Aware-GO VS Unaware-GO” contrast, since we are interested in investigating whether activity in motor areas is predicted by previous activity in LOC, when a motor response is required (i.e., in the GO condition). Thus, LOC will be used as seed ROI |  | **O3.2.1:** Significant predictive interactions between LOC and motor areas will suggest that, when the stimulus enters consciousness, awareness-related activity in LOC predicts subsequent activity in motor areas. This (expected) outcome will suggest that consciousness modulates subsequent response-related motor activity, by directly triggering activation of motor areas, as observed in Colombari et al., 2024<br>**O3.2.2:** If no significant interactions between LOC and MA will be highlighted, then it would mean that activity in motor areas is not predicted by LOC. Specifically, it could be surmised that in a *detection* task, consciousness does not modulate activation of motor areas, as observed in Colombari et al., 2024, where a |
|  |  |  | and later activity in motor areas will be investigated.<br>In addition to frequentist statistics, Bayesian statistics (with default prior) will be computed to test potential null effects<br>**Contrast to be computed:**<br>AWARE GO VS<br>UNAWARE GO<br>**ROI to be tested:**<br>LOC (as seed ROI)<br>Motor areas as predicted areas<br>**Time interval of interest:**<br>LOC: 150-350 ms after the stimulus onset (within this interval, about $\pm 4$ time points around the peak value -according to the peak waveform- will be selected)<br>MA: based on mean RTs, with a time window of $\pm 1.5$ sd around the mean |  | *discrimination* task was employed. The difference in the two tasks, indeed, consists in the type of motor response required: in the case of the discrimination task, the participant is asked to press one button or another according to the response. Conversely, in a detection task, the participant has to press a key when the target stimulus is detected. Thus, no selection of the response is needed. This difference could play a role in the relationship between consciousness and motor areas. |
| **Q4:** Does consciousness modulate activation of motor areas in ABSENCE of | **H4:** Consciousness modulates activation of motor areas, even if the motor response is not required | As Q3 | **A4.1:** A paired-sample one-tailed t-test will be computed in order to compare activity in Motor Areas in the NOGO condition. Thus, activity | As above | **O4.1.1:** A statistically significant t-test will suggest that, when a motor response is not provided, the inhibition required to withhold responding is stronger when the visual characteristic of the stimulus is consciously |
| motor response? | **Expected outcome:**<br>MA aware-NOGO>MA unaware-NOGO<br>LOC predicts MA (investigated by means of Granger Causality Analysis) |  | in Aware-NOGO and Unaware-NOGO trials will be contrasted.<br>In addition to frequentist statistics, Bayesian statistics (with default prior) will be computed to test potential null effects<br>**Contrast to be computed:**<br>AWARE NOGO VS UNAWARE NOGO<br>**ROI to be tested:**<br>Motor areas<br>**Time interval of interest:**<br>Based on mean RTs, with a time window of $\pm 1.5$ sd around the mean |  | perceived, compared to when no difference is perceived.<br>**O4.1.2:** If no difference between the tested conditions is observed, this will suggest that i) no inhibition is required to withhold responding, both in the Aware and Unaware condition, or ii) the inhibition is equally strong for the two conditions. |
|  |  |  | **A4.2:** With the aim of investigating the flow of activity occurring in the investigated brain areas also in the condition where no response is required, Granger Analysis will be performed on the “Aware-NOGO VS Unaware-NOGO” contrast. This will allow to investigate whether activity in motor areas is triggered by previous activity in LOC,<br>even when a motor response is not required. Thus, LOC will be used as seed ROI and later activity in motor areas will be investigated.<br>In addition to frequentist statistics, Bayesian statistics (with default prior) will be computed to test potential null effects<br>**Contrast to be computed:**<br>AWARE NOGO VS<br>UNAWARE NOGO<br>**ROI to be tested:**<br>LOC (as seed ROI)<br>Motor areas as predicted areas<br>**Time interval of interest:**<br>LOC: 150-350 ms after the stimulus onset (within this interval, about $\pm 4$ time points around the peak value -according to the peak waveform- will be selected)<br>MA: Based on mean RTs, with a time window of $\pm 1.5$ sd around the mean |  | **O4.2.1:** If significant predictive interactions between LOC and motor areas will be observed, then consciousness modulates subsequent activity in motor areas also in absence of a motor response. This could be due to inhibition of the response processes.<br>**O4.2.2:** If no significant interactions between LOC and MA will be highlighted, then LOC does not predict activity in motor areas in absence of motor response. |

## 4. Pilot study

In order to test the experimental paradigm, we pilot-tested the task.

A total of 10 right-handed participants (5 females and 5 males; mean age ± standard deviation: 21 years ± 1.0) took part in the pilot study. They all reported normal or corrected-to-normal vision and no history of neurological or psychiatric disorders. All of them provided written informed consent before starting the experiment.

After the first session, two participants dropped out the experiment, hence data from 8 participants were included in the statistical analyses. Moreover, in order to maintain an equal number of trials in both the conditions (i.e., Aware and Unaware), the percentage of Aware and Unaware trials was calculated and data from participants reporting a proportion of awareness equal or superior to 80% (i.e., 3 participants) were discarded from subsequent analysis. For this pilot study, we decided to raise the awareness threshold of acceptance to 80% (instead of 75%, that will be used in the experiment) in order to be more inclusive, given the low number of participants.

Thus, in total, data from 5 participants were included in the behavioral and functional analyses.

### 4.1 Preliminary Results

#### 4.1.1 Behavioral results

Raw data were processed by means of scripts created on Matlab (version R2017b; the MathWorks, Inc., Natick, MA). According to the participants’ responses, trials were sorted into the four experimental conditions (i.e., Aware-GO, Unaware-NOGO, Aware-NOGO and Unaware-GO). Aware trials were those trials in which the participant reported to perceive the thicker radius, while Unaware trials were those trials in which participants could not perceive that the radius was thicker. As specified in Section 2.8, trials with RTs lower than 150 ms or higher than 3SD from the mean were removed. After removal, we had on average 830.6 trials for the Aware-GO condition, 389.2 for the Unaware-NOGO condition, 738.8 trials for the Aware-NOGO condition and 491.4 for the Unaware-GO.

Subsequently, once assessed the normality of RTs and Awareness distributions (Shapiro-Wilk test. RTs distribution: W=0.824, p=0.125; Awareness distribution: W=0.817, p=0.112), the percentage of Awareness for the two conditions was calculated: in the GO condition, Aware trials represented on average 68.02% of the trials, while in the NOGO condition, Aware trials constituted the 59.82% of the trials. Paired sample (two-tailed) t-test performed with Jamovi (version 2.3.28) highlighted that there was no significant difference between the two conditions (t_(4)_ = 1.88, p = .134, Cohen’s d = .839). Similarly, mean RTs for Aware and Unaware trials in the GO condition were contrasted and the statistical analysis (Paired sample two-tailed t-test) revealed that mean RTs for the Aware condition (628.530 ms) and the Unaware condition (675.317 ms) were not statistically different (t(4) = – 1.77, p = .152, Cohen’s d = – .791). The behavioral results are depicted in Figure 3.

**Figure 3.**
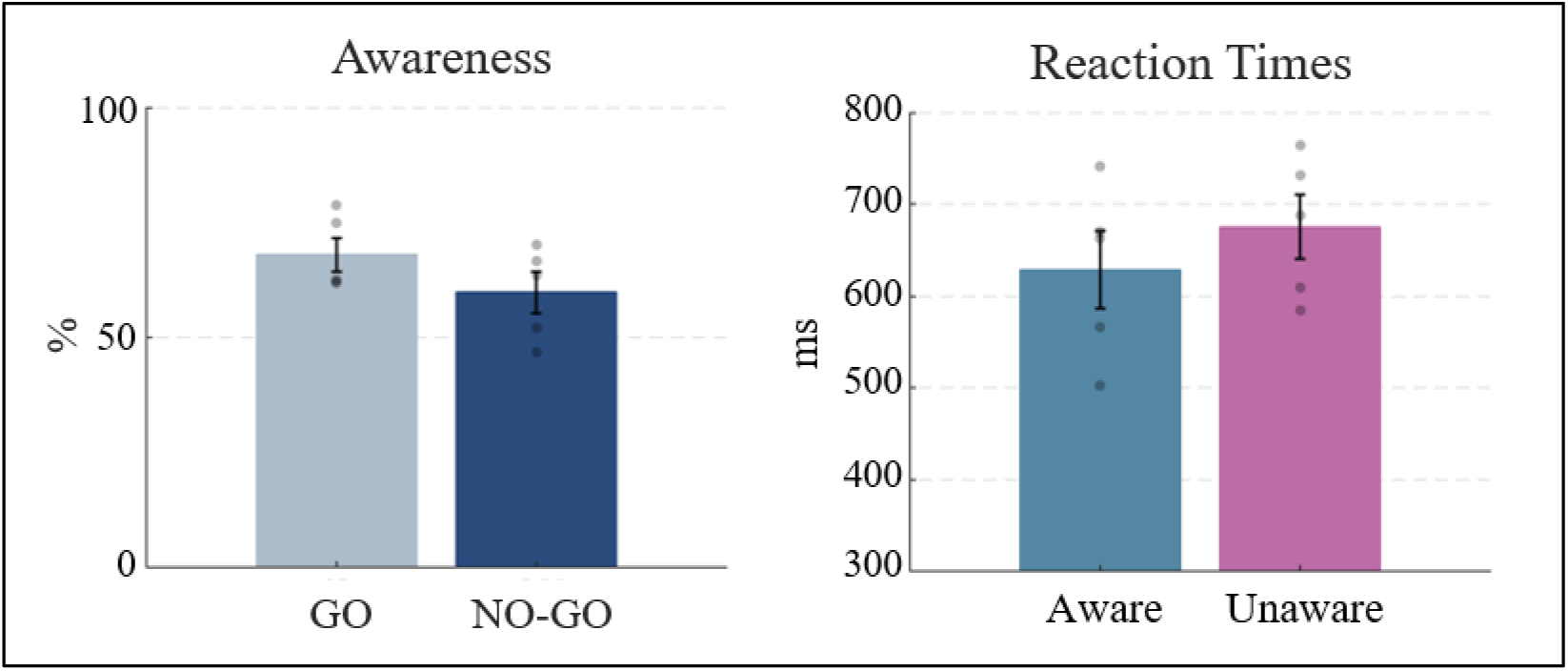
Behavioral results. The percentage of Awareness was calculated for both “GO” and “NOGO” conditions (on the left). Mean reaction times were calculated for Aware and Unaware trials only for the “GO” condition (on the right). No significant differences were observed. Error bars represent SEM and gray dots represent individual data points showing the data distribution.

Moreover, in order to verify that the employed paradigm works as planned and that participants performed the task accurately, analysis on catch trials was performed as described in Section 2.8.1 *Behavioral data.* As specified above, catch trials were those trials in which all the radii of the stimulus are equally thick. Hence, in those cases, participants should report not to see the thicker radius. As expected, they correctly reported not seeing the thicker radius on average the 96.47% of times (sd=2.49) in the Aware GO condition and the 98.36% (sd=1.89) in the Aware NOGO condition. Paired sample (two-tailed) t-test revealed no significant difference between the two conditions.

#### 4.1.2 EROS results

EROS data were pre-processed with a dedicated in-house software, P-POD (Pre-Processing of Optical Data, run in Matlab, version R2013b), as described in Section 2.8. Subsequently, we computed statistical analyses on pre-processed data by means of the dedicated in-house software package Opt-3d.

For this pilot study, participants’ individual structural MR images could not be acquired, so an estimated MR-based head model was individually created using the Softaxic Optic system (SofTaxic, E.M.S., Bologna, Italy) combined with a 3D optical digitizer (Polaris Vicra, NDI, Waterloo, Canada). EROS data were thus co-registered with the estimated MRI using a specific procedure performed in OCP software package (as specified above). Finally, co-registered data were transformed into MNI space for subsequent analyses.

For both GO and NOGO conditions, Aware and Unaware trials were contrasted. As shown in Figure 4, the Aware-GO vs Unaware-GO contrast replicated the results obtained by Colombari et al., 2024.

**Figure 4.**
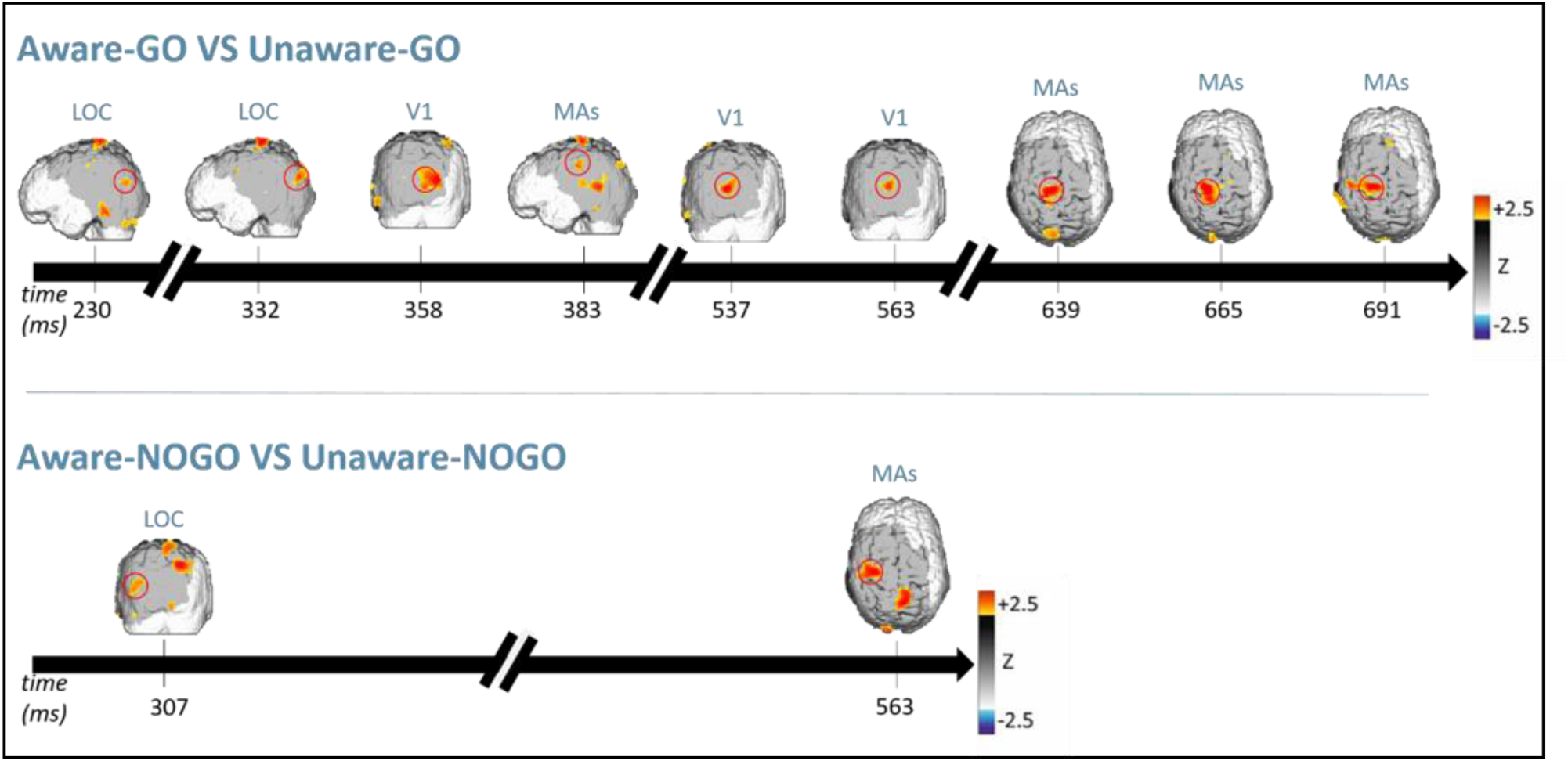
EROS results. Statistical parametric maps of the z-score difference computed contrasting Aware and Unaware trials in the GO (upper panel) and NOGO condition (lower panel). Each map represents a 25.6 ms interval.

In this contrast, indeed, we compared conditions in which the motor response was required, thus replicating the task carried out in the previously mentioned experiment. Also in this case, Aware trials elicited a sustained activation of LOC (230 and 332 ms after the stimulus onset), followed by the recurrent activation of the primary visual cortex (V1) and the motor areas (MA) at later stages of stimulus processing.

Similarly, contrasting Aware and Unaware trials in the condition where the motor response was not required (i.e., the NOGO condition), greater activation of LOC was elicited in a timing comparable to that of the contrast just mentioned above (i.e., 307 ms after the stimulus presentation). Interestingly, also in this case awareness-related processing elicited activity in the motor areas, 563 ms after the stimulus onset, despite in this condition no response was required, possibly suggesting an inhibition to respond for the NOGO trials.

### 4.2 Preliminary Discussion

The aim of the present pilot study was to assess whether the task and the experimental procedure were suitable to investigate the study’s research questions.

As described in Section 4.1, the pilot study successfully replicated the trend of activations observed by Colombari et al., 2024, suggesting that the proposed study proves to be feasible in terms of methodology. For the sake of clarity, it is important to point out that the preliminary results reported here do not reach the statistical level of significance. This outcome was expected as data from only 5 participants were included in the analysis. For the same reason, we decided not to perform Granger Causality analysis as for this kind of analysis results from 5 participants would have been uninformative. Nevertheless, it was possible to observe that the proposed task could elicit a pattern of activation similar to that observed by Colombari et al., 2024, suggesting that the experimental paradigm proposed to investigate the research questions is suitable.

## Data availability

Upon acceptance of the Stage 2 registered report, we will share all raw and processed anonymized data as well as study materials publicly available as open data. Pilot raw and processed data can be found on this link: https://osf.io/ebfu3/?view_only=9ec2e6bf32ba4a8bb8b858639ec40a59

## Code availability

All analysis codes will be made publicly available upon acceptance of the Stage 2 registered report.

## Acknowledgments

The present project is supported by the grant program “Funding Consciousness Research with Registered Reports” and Fondazione Cassa di Risparmio di Verona, Vicenza, Belluno e Ancona “Ricerca scientifica d’eccellenza 2018”, “Emergence of Consciousness: From neural dynamics to complex conscious behaviour” grant no. 2018.0861; SS is supported by #NEXTGENERATIONEU (NGEU) and funded by the Ministry of University and Research (MUR), National Recovery and Resilience Plan (NRRP), project MNESYS (PE0000006)—“A Multiscale integrated approach to the study of the nervous system in health and disease” (DN. 1553 11.10.2022), CM is supported by MIUR D.M. 737/2021—“Neural correlates of perceptual awareness: from neural architecture to the preservation of conscious vision in brain tumor patients”.

## Author contributions

**EC** Conceptualization, Methodology, Software, Validation, Formal analysis, Investigation, Data Curation, Writing – Original Draft, Visualization, Funding Acquisition; **GP** Methodology, Formal Analysis, Investigation, Data Curation, Writing – Review & Editing; **SM** Methodology, Investigation, Writing – Review & Editing **CM** Methodology, Software, Data Curation, Writing – Review & Editing, Supervision; **SS** Conceptualization, Methodology, Resources, Writing – Review & Editing, Supervision, Project administration, Funding acquisition.

## Competing interests

The authors declare no competing interests.

## Notes

https://osf.io/ebfu3/

### Competing Interest Statement

The authors have declared no competing interest.

### Summary of Updates

The study design table has been improved

