## Supplementary Table 1 for "The role of extra-striate areas in conscious motor behavior: a registered report with Fast-Optical Imaging" (from another version of this preprint: version 3, 2024-12-12)

| Title | Authors | Journal | Year of publication | Sample size |
|---|---|---|---|---|
| Shades of gray matter: Noninvasive optical images of human brain responses during visual stimulation | Gratton, G., Corballis, P.M., Cho, E., Fabiani, M., & Hood, D.C. | Psychophysiology | 1995 | 3 |
| Fast and Localized Event-Related Optical Signals (EROS) in the Human Occipital Cortex: Comparisons with the Visual Evoked Potential and fMRI | Gratton, G., Fabiani, M., Corballis, P.M., Hood, D.C., Goodman-Wood, M.R., Hirsch, J., Kim, K., Friedman, D., & Gratton, E. | Neuroimage | 1997 | 3 |
| Toward noninvasive 3-D imaging of the time course of cortical activity: Investigation of the depth of the Event-Related Optical Signal | Gratton, G., Sarno, A., Maclin, E., Corballis, P.M., and Fabiani, M. | Neuroimage | 2000 | 9 |
| Comparison of Neuronal and Hemodynamic Measures of the Brain Response to Visual Stimulation: An Optical Imaging Study | Gratton, G., Goodman-Wood, M.R., & Fabiani, M. | Human Brain Mapping | 2001 | 8 |
| The event-related optical signal (EROS) in visual cortex: Replicability, consistency, localization, and resolution | Gratton, G. & Fabiani, M. | Psychophysiology | 2003 | 8 |
| Fast cerebral functional signal in the 100-ms range detected in the visual cortex by frequency-domain near-infrared spectrophotometry | Wolf, M., Wolf, U., Choi, J.H., Toronov, V., Paunescu, L.A., Michalos, A., & Gratton, E. | Psychophysiology | 2003 | 4 |
| Fast optical imaging of frontal cortex during active and passive oddball tasks | Low, K.A., Leaver, E., Kramer, A.F., Fabiani, M., & Gratton, G. | Psychophysiology | 2006 | 18 |
| Preattentive Change Detection Using the Event-Related Optical Signal | Tse, C.Y. & Penney, T.B. | IEEE Engineering In Medicine And Biology Magazine | 2007 | 19 |
| Imaging cortical dynamics of language processingwith the event-related optical signal | Tse C.Y., Lee C.L., Sullivan J., Garnsey S.M., Dell G.S., Fabiani M., and Gratton G. | Proceedings of the National Academy of Sciences | 2007 | 16 |
| Event-related fast optical signal in a rapid object recognition task: Improving detection by the independent component analysis | Medvedev, A. V., Kainerstorfer, J., Borisov, S. | Brain research | 2008 | 10 |

| | | | | |
|---|---|---|---|---|
| | V., Barbour, R.L., & VanMeter, J. | | | |
| "Seeing" electroencephalogram through the skull: imaging prefrontal cortex with fast optical signal | Medvedev, A. V., Kainerstorfer, J.M., Borisov, S. V., Gandjbakhche, A.H., & VanMeter, J. | Journal of Biomedical Optics | 2010 | 11 |
| Frontoparietal Traffic Signals: A Fast Optical Imaging Study of Preparatory Dynamics in Response Mode Switching | Baniqued, P., L.; Low, K.,A.; Fabiani, M; Gratton, G. | Journal of Cognitive Neuroscience | 2013 | 15 |
| The time-course of cortical responses to speech revealed by fast optical imaging | Toscano, J.C., Anderson, N.D., Fabiani, M., Gratton, G., & Garnsey, S.M. | Brain and Language | 2018 | 12 |
| Quantifying fast optical signal and event-related potential relationships during a visual oddball task | Proulx, N., Samadani, A.A., & Chau, T. | Neuroimage | 2018 | 15 |
| Spatiotemporal dynamics of attentional orienting and reorienting revealed by fast optical imaging in occipital and parietal cortices | Parisi, G., Mazzi, C., Colombari, E., Chiarelli, A.M., Metzger, B.A., Marzi, C.A., & Savazzi, S. | Neuroimage | 2020 | 18 |
| Fronto-occipital mismatch responses in pre-attentive detection of visual changes: Implication on a generic brain network underlying Mismatch Negativity (MMN) | Tse, C. Y., Shum, Y. H., Xiao, X. Z., Wang, Y. | Neuroimage | 2021 | 22 |
| Frontotemporal network in automatic / pre-attentive detection of abstract change: An event-related optical signal (EROS) study | Xiao X.Z, Wang Y. , Wong G.C.S., Zhao K., Tse C.Y. | Neuropsychologia | 2022 | 30 |
| Bypassing input to V1 in visual awareness: A TMS-EROS investigation | Knight, R.S., Chen, T., Gratton, G., Fabiani, M., Savazzi, S., Mazzi, C., & Beck, D.M. | Neuropsychologia | 2024 | 12 |
| | | | mean | 12.944 |
| | | | SD | 7.008 |
